# Engineering Microbial Physiology with Synthetic Polymers: Cationic Polymers Induce Biofilm Formation in *Vibrio cholerae* and Downregulate the Expression of Virulence Genes

**DOI:** 10.1101/066563

**Authors:** Nicolas Perez-Soto, Lauren Moule, Daniel N. Crisan, Ignacio Insua, Leanne M. Taylor-Smith, Kerstin Voelz, Francisco Fernandez-Trillo, Anne Marie Krachler

## Abstract

Here we report the first application of non-bactericidal synthetic polymers to modulate the physiology of a bacterial pathogen. Poly(*N*-[3-(dimethylamino)propyl]methacrylamide) (**P1**) and poly(*N*-(3-aminopropyl) methacrylamide) (**P2**), cationic polymers that bind to the surface of *V. cholerae,* the infectious agent causing cholera disease, can sequester the pathogen into clusters. Upon clustering, *V. cholerae* transitions to a sessile lifestyle, characterised by increased biofilm production and the repression of key virulence factors such as the cholera toxin (CTX). Moreover, clustering the pathogen results in the minimisation of adherence and toxicity to intestinal epithelial cells. Our results suggest that the reduction in toxicity is associated with the reduction to the number of free bacteria, but also the downregulation of toxin production. Finally we demonstrate that these polymers can reduce colonisation of zebrafish larvae upon ingestion of water contaminated with *V. cholerae*. Overall, our results suggest that the physiology of this pathogen can be modulated without the need to genetically manipulate the microorganism and that this modulation is an off-target effect that results from the intrinsic ability of the pathogen to sense and adapt to its environment. We believe these findings pave the way towards a better understanding of the interactions between pathogenic bacteria and polymeric materials and will underpin the development of novel antimicrobial polymers.

## Introduction

Infectious diseases are one of the greatest medical challenges today, because of the increase in occurrence of antimicrobial resistance in bacterial infections.^1,2^ Thus, new approaches to tackle these diseases are needed. In particular novel approaches that can replace or enhance current antimicrobials are highly desired.^3–5^ Polymeric materials have often been postulated as alternatives in these areas, either as delivery vehicles for antimicrobials,^6,7^ or as novel antimicrobial polymers,^8–11^ and materials that can interfere with microbial adhesion.^12–15^ Polymeric materials are especially attractive in these applications because of their multivalency, ease of manufacturing and the potential to precisely control polymer length and composition.^16^

Despite this vast progress in polymer technology, and the amount of “antimicrobial” polymeric structures described to date, little is known about the effect of these materials on the physiology of the targeted pathogens. However, the ability of pathogenic bacteria to sense and respond to changes in their environment is responsible for their ability to adapt to different lifestyles, and colonise different niches, including new hosts. For instance, the transition from free-living to an adherent state in bacteria is often mediated by a combination of chemical and physical cues. This transition leads to changes in gene expression, and in microbes with pathogenic potential it can result in the upregulation of virulence genes. At present, our understanding of how polymeric materials may be triggering these complex chemical and physical cues is very limited, and this lack of understanding is compromising our ability to develop these materials.

As a first approach towards this understanding, in this manuscript we report the effect of poly(*N*-[3-(dimethylamino)propyl]methacrylamide) (**P1**) and poly(*N*-(3-aminopropyl)methacryla-mide) (**P2**) on the human pathogen *Vibrio cholerae*. We demonstrate that cationic polymeric materials that can sequester this pathogen into clusters and thus have the potential to inhibit its adhesion to the host, result in a complex modulation of virulence factors and gene expression. While biofilm production is increased, other virulence factors are suppressed, including the main causative agent of the cholera disease, the cholera toxin. Finally, we demonstrate that these polymeric materials can indeed inhibit the colonisation by *V. cholerae* of relevant *in vitro* and *in vivo* models.

## RESULTS AND DISCUSSION

*V. cholerae* is a Gram-negative bacterium responsible for several million incidences of gastrointestinal disease and up to 142,000 deaths every year.^17,18^ Within the human host, the bacterium initiates a virulence programme including the induction of colonisation factors and toxins, in response to the chemical and physical cues experienced during colonisation and adhesion. In aquatic environments, *V. cholerae* persists by forming biofilms on the surfaces of phytoplankton, zooplankton and chitin debris.^19,20^ Biofilms offer a protective environment both against aquatic predators or in the host environment. Overall, the ability to switch between motile and biofilm lifestyles, along with the carefully controlled induction of virulence factors, is central to the establishment of disease and the emergence of cholera epidemics.^21,22^

**Scheme 1.**
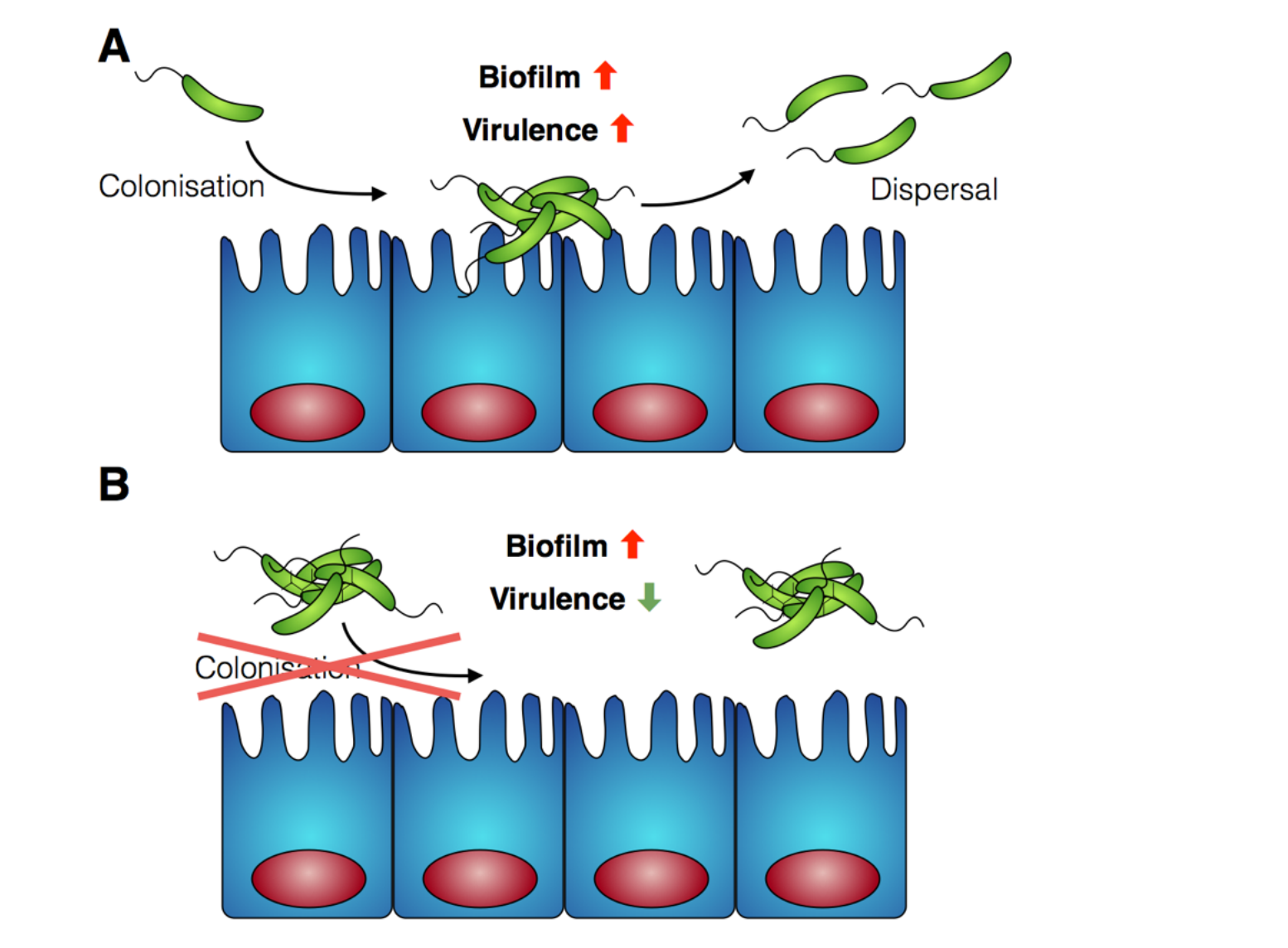
Schematic representation of *V. cholerae* infection and the effect of the polymers reported in this work. **A**) *V. cholerae* infection is associated with the colonisation of the gut epithelia and the upregulation of biofilm formation and virulence. **B**) In the presence of the polymers reported herein, colonisation is minimised by sequestering the pathogen in clusters. Additionally, a repression of virulence is observed.

Cholera is a particular concern in environments where there is poor sanitation and thus, access to clean water is not available. Therefore, efforts have been made to develop prophylactic approaches to the treatment of cholera, including the development of vaccines,^23^ or filtration devices to clean water. Most of these filtration devices simply physically restrict the passage of *V. cholerae* (and other pathogens) through the pores of a membrane, although recent efforst are being made to develop polymeric materials that can selectively bind to microbial strains.^24,25^ Based on our previous work with closely related microorganism Vibrio harveyi,^26,27^ we decided to investigate what would be the response of *V. cholerae* to polymeric materials designed to bind to its surface and sequester the pathogen into clusters. Previously reported microbial sequestrant poly(*N*-[3-(dimethylamino)propyl]methacrylamide) (**P1**)^27,28^ was synthetised via free radical polymerisation, using 2-mercaptoethanol as a chain-transfer agent (see ESI: section 4.1, **Fig. S1**^†^ for experimental details and characterisation). This polymer carrying tertiary amine residues should be mainly protonated at physiological pH providing an overall polycationic charge.^28,29^ We anticipated that this cationic polymer should therefore bind to the surface of negatively charged *V. cholerae*, and sequester the microorganism into clusters. Optical microscopy revealed that cluster formation proceeded via initial nucleation of small layers or sheets of bacteria, which increased in size both by lateral interaction with additional bacteria, as well as stacking of bacteria on existing sheets to form clusters over the first 15 minutes, and then remained stable over the duration of the experiment (Fig. 1A and B). Polymer-induced bacterial clusters were stable and had an extended three-dimensional structure (Fig. 1C and Fig. S3). This capture of bacteria in polymer clusters also compromised motility, as evidenced by the reduced migration in soft agar plates of *V. cholerae* in the presence of **P1** (Fig. S4). This effect was dose dependent, with lower motility observed for the higher concentration of polymer. Full motility was restored upon exposure of clusters to high salt concentration (Fig. S4, bottom), confirming that clustering was driven by electrostatic interactions that could be easily disrupted in the presence of competing electrolytes.

**Fig. 1.**
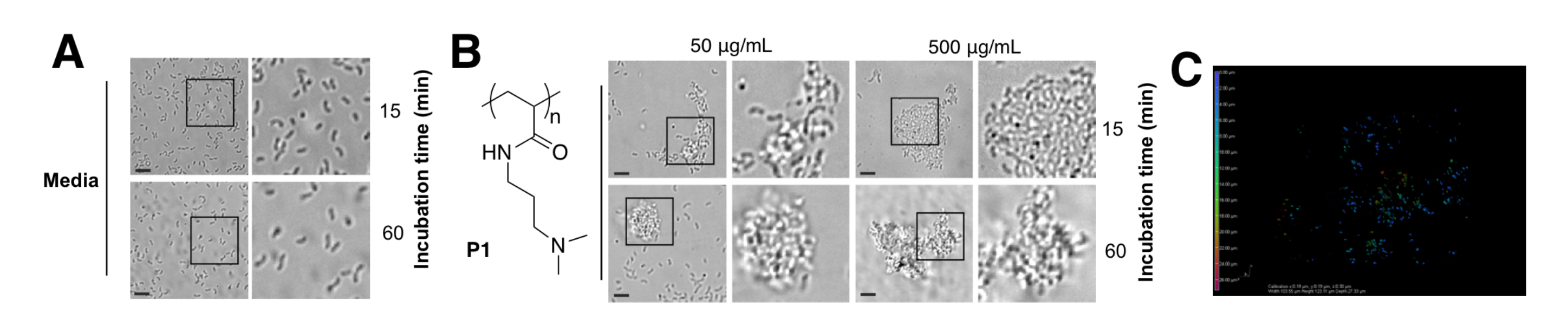
Optical micrographs of *V. cholerae* N16961 suspensions incubated in PBS (pH 7.4) in the absence (A) and in the presence of **P1** (B) at a polymer concentration of 50 or 500 μg/mL for 15 or 60 min. Area within the square has been expanded (right row) for clarity. C) Spinning disk fluorescent micrograph of GFP-*V. cholerae* N16967 incubated in PBS (pH 7.4) in the presence of **P1** at a polymer concentration of 500 μg/mL for 15 min. Color coding represents z-depth.

Having identified that **P1** was able to sequester *V. cholerae* into clusters, we then evaluated the viability of the pathogen in the presence of this polymer. Cationic polymers are commonly reported as bactericidal materials,^8–11^ although small changes in structure and dose can result in significant differences in activity and toxicity. This effect is often microbe specific, and in particular **P1** has been reported to have a minimum inhibitory concentration of 10-25 μg/mL for Escherichia coli,^30^ while showing no effect on V. harveyi’s viability and growth at concentations as high as 500 μg/mL.^27^ Viability in *V. cholerae* was therefore analysed by monitoring the effect of **P1** in bacterial proliferation and viability under physiological conditions. Bacterial proliferation is often measured by monitoring the effect of a substance over the optical density at 600 nm (OD_600_) of a culture of bacteria, since the optical density of the sample can be correlated to the number of bacterial cells in the culture.^31^ However, OD_600_ can be affected by clustering. The presence of clusters can result in an increase in the optical density of the culture due to the bigger size and slower disfussion of these clusters, or a reduction of OD_600_, due to coaccervation of the clusters once they reach a critical size.^24^ Due to this limitation, we monitored production of green fluorescent protein (GFP) as a proxy of bacterial growth. Co-incubation of GFP expressing *V. cholerae* with different concentrations of **P1** was followed by monitoring GFP fluorescence over 25 hours which showed that bacterial proliferation was not significantly affected (Fig. 2A). However, it was unclear from these experiments whether there could be cellular damage commonly observed with highly charged cationic polymers. Flow cytometry of bacterial samples exposed to polymers and LIVE/DEAD® cell viability stains allowed us to determine membrane integrity of *V. cholerae* sequestered in clusters at the experimental endpoint (Fig. 2B). Again, membrane integrity was largely unaffected even following overnight incubation.

**Fig. 2.**
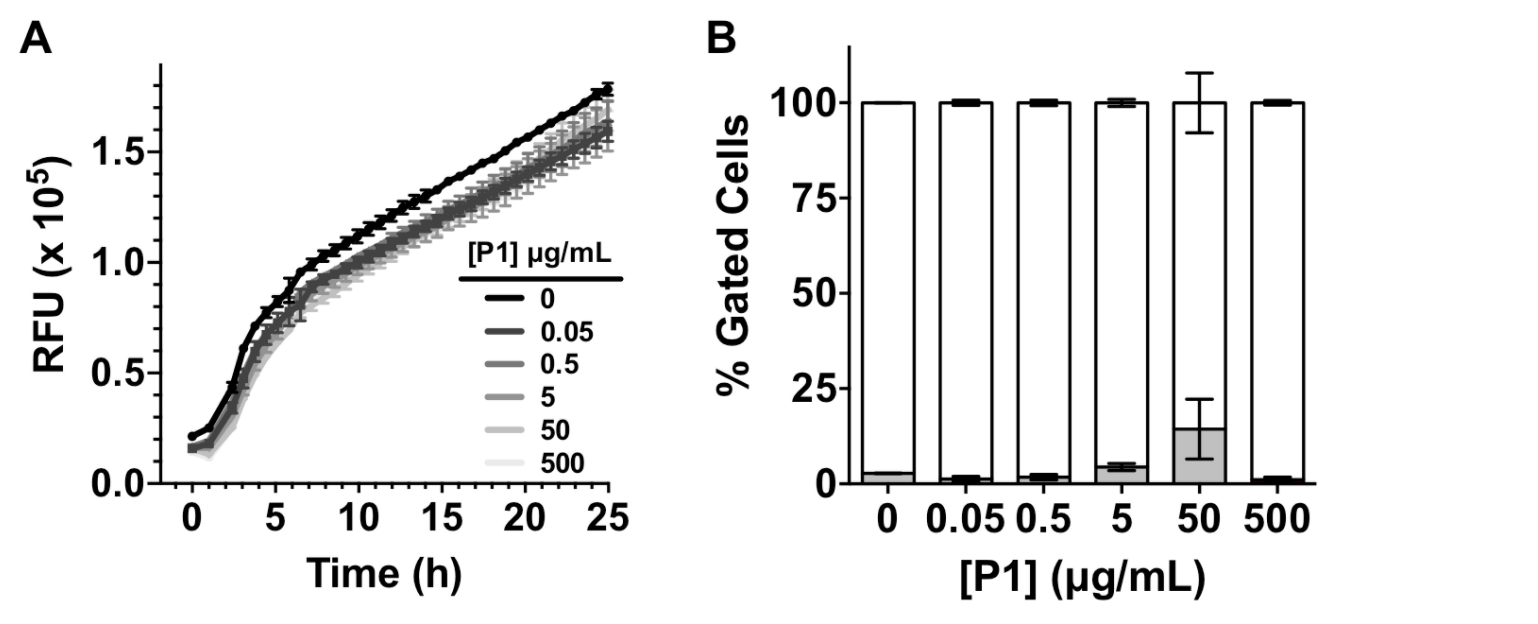
A) Relative Fluorescent Units (RFU) for GFP expressing *V. cholerae* N16961 in the absence and presence of increasing amounts of **P1**. Initial OD_600_ = 0.2. Results are means ± s.e.m. of three independent experiments. B) Normalised population of *V. cholerae* N16961 incubated in the absence and presence of **P1**. Normalised population is presented as the percentage of green (hollow bars, non-permeable/viable) and red (grey bars, high permeability/dead) cells as measured by flow cytometry. Bacteria were treated with i-PrOH as a negative control. Initial OD_600_ = 1. Results are means ± s.e.m. of three independent experiments.

Having identified that **P1** was non-toxic to *V. cholerae*, and that it was able to induce clustering of this pathogen, we then decided to investigate whether the physiology of the pathogen was being affected following this clustering. Visual inspection by optical microscopy suggested similarities between *V. cholerae* biofilms and these polymer-induced clusters (Fig. 1). Interestingly, infectious *V. cholerae* are often taken up as small biofilms, from which bacteria escape to colonise the epithelium. Once bound to host cells, bacteria initiate microcolony formation, before eventually exiting the host’s gastrointestinal tract, often following re-organisation into biofilms,^32,33^ to cause environmental dispersal and onward-transmission (Scheme 1A). The ability to transition between motile and sessile states is thus key to *V. cholerae*’s virulence regulation. With our polymers bacterial motility was largely abolished, by physical deposition into polymer-based clusters. We therefore investigated whether polymer-induced clustering would also affect *in vitro* biofilm formation. *V. cholerae* biofilms are mainly composed of *Vibrio* exopolysaccharide (VPS) ^22,34^ which can be stained using crystal violet,^35,36^ a chromogenic probe that interacts with negatively charged biopolymers. UV-Vis analysis of these polymer-induced clusters revealed an increased retention of the dye in the presence of **P1**, suggesting higher levels of extracellular exopolysaccharides (Fig. 3A). Additionally, staining of these cultures with Hoechst revealed increased amounts of extracellular DNA (eDNA) in the presence of **P1** (Fig. 3C). eDNA is another common component of *V. cholerae* biofilms,^22^ and in our case, patches of diffuse staining could be seen around the polymer-induced clusters. In an attempt to quantify the levels of eDNA, the intensity of the blue channel was measured (**Fig. S6A**). We observed an increase in the intensity of this blue channel for some of the samples incubated with **P1**, although the variablity in the samples prevented quantification of this increase. Interestingly, treatment of polymer-induced biofilms with DNAse I abolished the additional blue fluorescence, confirming the released substance was indeed eDNA (**Fig. S6A**, grey dots).

**Fig. 3.**
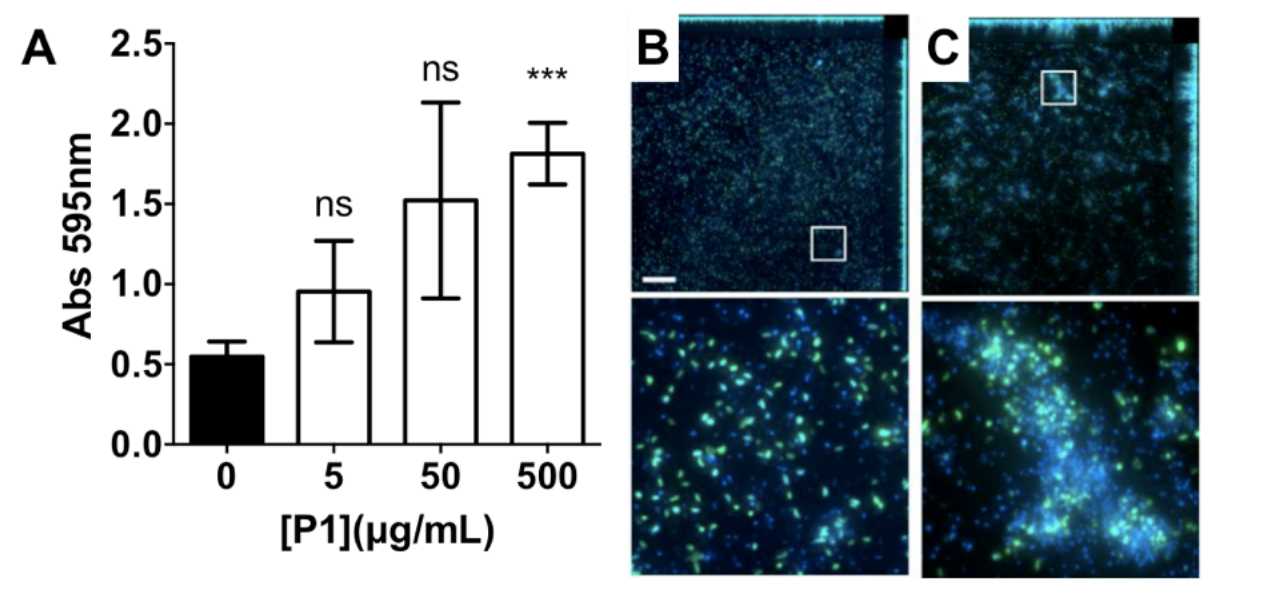
A) Absorbance at 595 nm for *V. cholerae* N16961 cultures in the absence and presence of **P1**. Analysis of variance (ANOVA), followed by Tukey’ s post hoc fest, was used to test for significance. Statistical significance was defined as p<0.05 (*), p<0.01 (**) or p<0.001 (***). Confocal fluorescence micrographs of GFP expressing *V. cholerae* N16961 cultures in the absence (B) and presence of 500 μg/mL **P1** (C). Scale bar, 50 μm. Area within the square has been expanded (bottom row) for clarity.

In the environment, transition of *V. cholerae* from planktonic to biofilm-associated growth leads to suppression of virulence-specific genes. Since sequestration into polymer-induced clusters promoted a sessile state, similar to biofilm-associated growth, we investigated what impact this clustering would have on the transcriptional regulation of key virulence factors. We were particularly interested in the regulation of cholera toxin (CTX), which is the main causative agent of cholera disease. For this purpose, we created a series of transcriptional reporter strains, by introducing, via conjugation, a variant of the pRW50 plasmid containing oriT into *V. cholerae* (Fig. 4A). Thus, we could follow transcription from *V. cholerae* promoters using β-galactosidase assays. An infection model with Caco-2 intestinal epithelial cells was used because contact with cultured epithelial cells has been described as a strong activating cue for expression of virulence factors.^37^

**Fig. 4.**
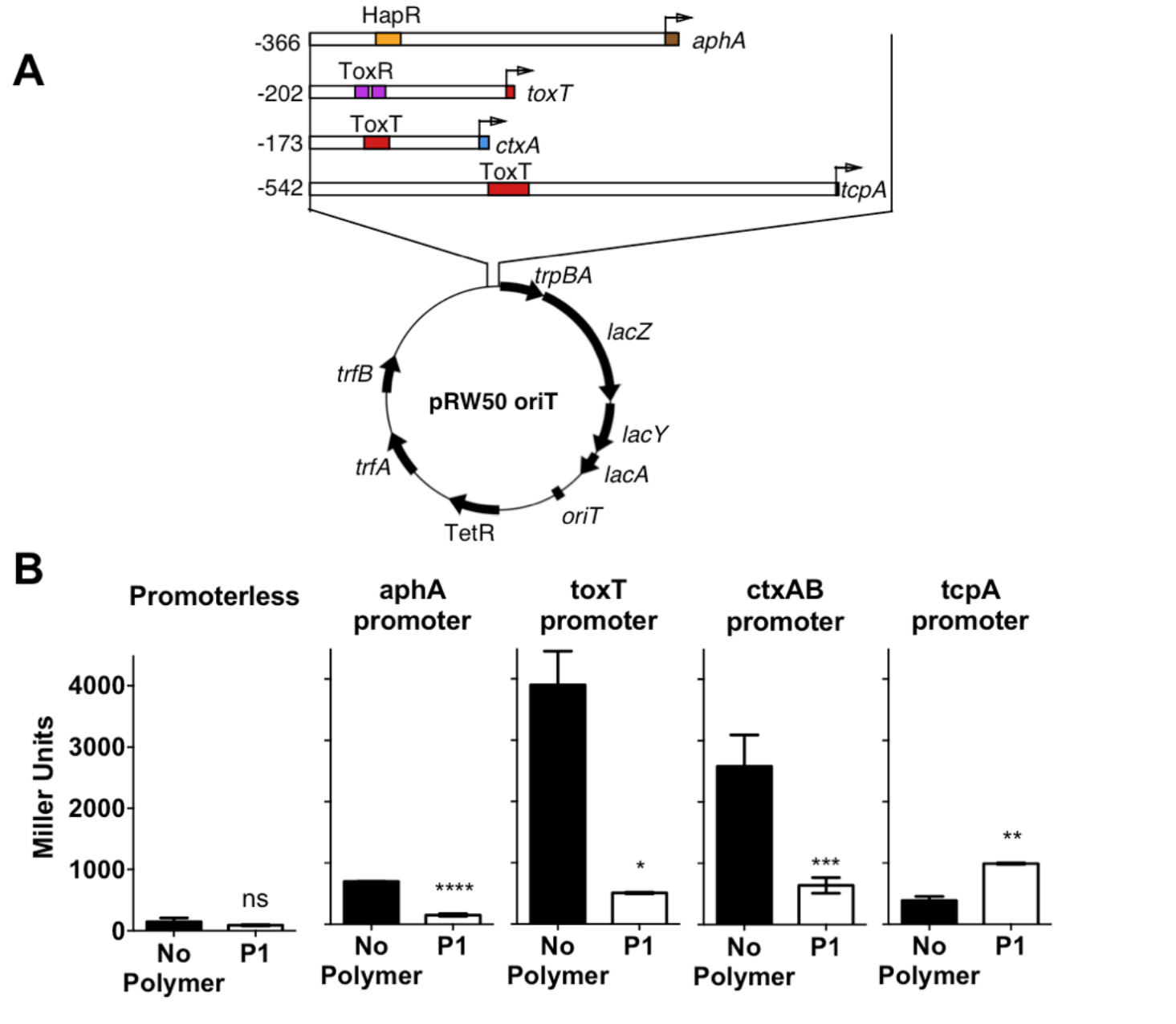
A) Schematic depiction of the reporter plasmid (pRW50-*oriT*) and promoter regions of *aphA*, *toxT*, *ctxA* and *tcpA* cloned as transcriptional fusions to *lacZ* in pRW50-oriT. Numbering refers to base number relative to the transcriptional start site and promoter binding sites for HapR, ToxR and ToxT are indicated. B) Promoter activities of *aphA-*, *toxT*-, ctxAB- and tcpA-lacZ fusions following infection of Caco-2 cells for 7 hours in the absence (black) or presence of 500 μg/mL **P1** (hollow). Student’s paired t-test was used to test for significance. Statistical significance was defined as p<0.05 (*), p<0.01 (**), p>0.001 (***), or p>0.0001 (****).

The viability of this cell line in the presence of the polymer was assessed first, using a lactate dehydrogenase release (LDH) assay to probe cellular membrane integrity (**Fig. S9**). Not suprisingly, **P1** compromised membrane integrity of the mammalian cells at the highest concentrations tested (≥ 50 μg/mL), in agreement with the reported toxicity of cationic materials.^8–11^ However, we anticipated that this toxicity could be reduced following incubation with bacteria, as the cationic charge will not be exposed following interaction with the pathogen’s negatively charged membrane. This was indeed the case when we evaluated the potential of this polymer to inhibit colonisation by *V. cholerae* of this intestinal epithelial cell line (Fig. 6C). No additional toxicity was observed when compared to that of *V. cholerae* alone, suggesting that incubation of **P1** with the bacteria resulted in a reduction of this polymer’s toxicity.

Then, we investigated expression of a series of regulators of virulence (Fig. 4A). AphA is a master regulator of virulence that is required for the activation of *tcpP*.^38^ TcpP, in turn, activates *toxT*, which activates downstream virulence regulators, including those responsible for the production of cholera toxin (*ctxAB*) and the toxin-coregulated pilus (*tcpA*). In our experiments, *V. cholerae* was first incubated with the polymer, to induce clustering, and these clusters where then used to colonise Caco-2 monolayers. Cells were then incubated for 7 hours, a time we determined to yield a maximum level of adherence and cytotoxicity.^39^ It was evident from these experiments that both *toxT* and *ctxAB* promoters were significantly repressed in the presence of **P1**, with up to 85% and 75% reduction respectively (Fig. 4B). Overall, neither *aphA* and *tcpA* were significantly activated, when compared to a promoterless strain.

Repression of *ctxAB* was very intriguing because *V. cholerae* uses quorum sensing regulation to repress virulence factor production and biofilm formation at high cell densities.^40,41^ However, virulence regulation is a complex phenomenum, where not only quorum sensing is involved, but complex chemical and physical cues are integrated to determine phenotypic expression.^42,43^ Based on this we decided to investigate whether this regulation was unique for **P1**, or other cationic polymer could trigger the same response in *V. cholerae*. Primary amine containing polymer **P2**^30^ (Fig. 5) was thus prepared from commercially available *N*-(3-aminopropyl)methacrylamide hydrochloride, using a similar experimental procedure (see ESI: section 4.2, **Fig. S2**† for experimental details and characterisation). Overall, the effect of **P2** in *V. cholerae* was very similar to that of **P1**. Clustering (Fig. 5A-C), reduced motility (**Fig. S4**) and a similar modulation of virulence was observed (Fig. 5E-H), although the presence of a primary amine in this polymer made it more toxic to both the pathogen (Fig. 5D and **Fig. S3**) and the host cells (**Fig. S9**). Thus, for functional experiments, we focused on investigating the effects of the highest effective, non-toxic to *V. cholerae* concentration of both polymers, *i.e.* 500 μg/mL for **P1** and 50 μg/mL for **P2**.

Since sequestration of *V. cholerae* into polymer-induced clusters repressed the transcription of key virulence factors during infection of Caco-2 cells, we asked what impact this clustering would have on the colonisation and toxicity by *V. cholerae* to these intestinal epithelial cells. In principle, sequestration should minimise the amount of free pathogen, thus minimising its potential to colonise this cell line. Repression of CTX under these conditions should give the additional benefit of reducing the toxicity of this bacteria to the cells. As expected, incubation of *V. cholerae* with both polymers prior to infection led to a significant reduction in bacterial attachment to host cells, as determined by dilution plating/CFU counts, and the effect of **P2** on this colonisation was much more pronounced than for **P1** (Fig. 6A). Imaging of Caco-2 cells infected with either planktonic (Fig. 6B, No polymer) or clustered *V. cholerae* (Fig. 6B, 5-500 μg/mL) also revealed a decrease in *V. cholerae* mediated toxicity as a result of bacterial clustering. Host cells infected with clustered bacteria showed more intact cell-cell junctions than cells infected with planktonic bacteria, in particular at low concentrations where no significant toxicity of the polymers to Caco-2 cells was observed (**Fig. S9**). This protective effect of the polymers was also observed when cytotoxicity was measured by LDH release assays, following infection with planktonic (Fig. 6C, 0 μg/mL) or clustered *V. cholerae* (Fig. 6C, 0.05-500 μg/mL). As described before, there was a cumulative effect, and the toxicity of polymer treated bacteria was lower than that of the individual components. This observation is in agreement with polymers sequestering bacteria and minimising the pathogenic burden, but conversely, bacteria “sequestering” the polymers and minimising the amount of cationic moieties exposed. Thus, the overall toxicity is reduced below that of the individual compontent. Additionally, we tested the specific effect of clustering on CTX activity, by measuring cAMP activity in infected cells (Fig. 6D). Elevated cAMP production due to CTX activity is the main hallmark of cholera infection, and is responsible for diarrhea in humans. While cAMP levels were significantly elevated in *V. cholerae* infected Caco-2 cells (Fig. 6D, 0 μg/mL), clustering of bacteria with both polymers decreased cAMP levels during infection, in agreement with the reduction in ctxAB transcription observed before (Fig. 4B and Fig. 5H).

**Fig. 5.**
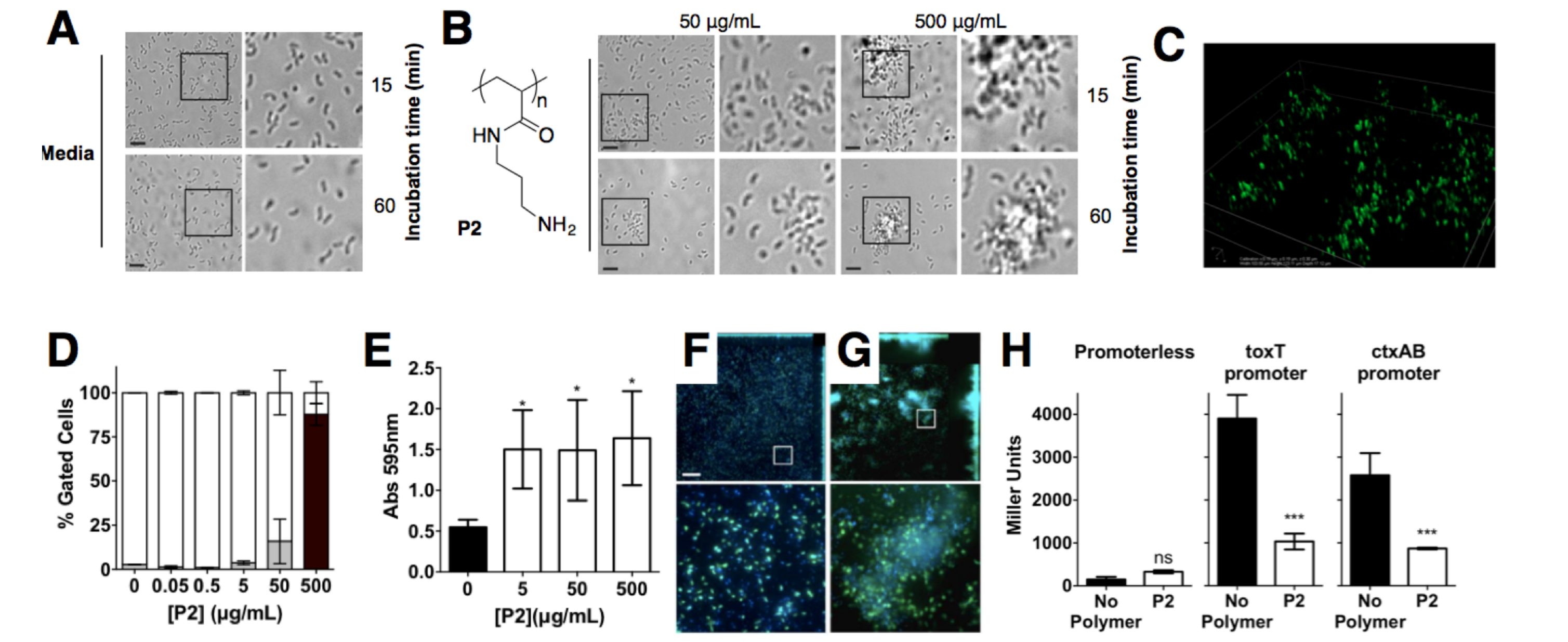
Optical micrographs of *V. cholerae* N16961 suspensions incubated in PBS (pH 7.4) in the absence (A) and in the presence of **P2** (B) at a polymer concentration of 50 or 500 μg/mL for 15 or 60 min. Area within the square has been expanded (right row) for clarity. C) Spinning disk fluorescent microscopy of GFP-*V. cholerae* N16961 incubated in PBS (pH 7.4) in the presence of **P2** at a polymer concentration of 500 μg/mL for 15 min. D) Normalised population of *V. cholerae* N16961 incubated in the absence and presence of **P2**. Normalised population is presented as the percentage of green (hollow bars, non-permeable/viable) and red (grey bars, high permeability/dead) cells as measured by flow cytometry. Bacteria were treated with i-PrOH as a negative control. Initial OD_600_ = 1. Results are means ± s.e.m. of three independent experiments. E) Absorbance at 595 nm for *V. cholerae* N16961 cultures in the absence and presence of **P2**. Analysis of variance (ANOVA), followed by Tukey’s post hoc test, was used to test for significance. Statistical significance was defined as p<0.05 (*), p<0.01 (**) or p<0.001 (***). Confocal fluorescence micrographs of GFP expressing *V. cholerae* N16961 cultures in the absence (F) and presence of 50 μg/mL **P2** (G). Scale bar, 50 μm. Area within the square has been expanded (bottom row) for clarity. H) Promoter activities of *toxT-* and *ctxAB*-*lacZ* fusions following infection of Caco-2 cells for 7 hours in the absence (black) or presence of 50 μg/mL **P2** (hollow). Student’s paired t-test was used to test for significance. Statistical significance was defined as p<0.05 (*) or p<0.001 (***).

**Fig. 6.**
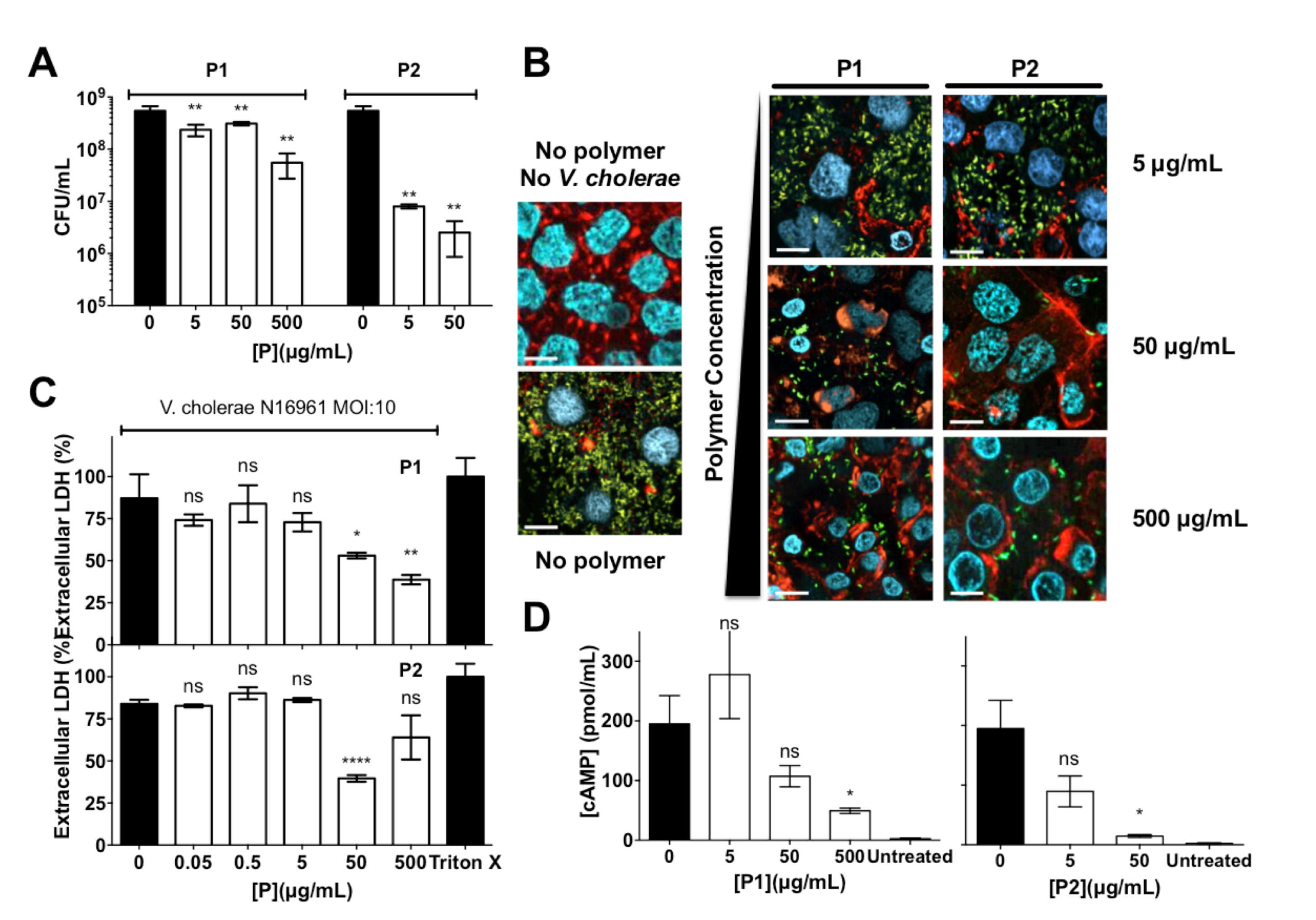
**A)** Number of colony forming units per mL (CFU/mL) of GFP expressing *V. cholerae* N16961 from Caco-2 cells incubated in the absence and presence of polymer treated GFP expressing *V. cholerae* N16961 cultures, following washing and lysis of Caco-2 cells. **B**) Confocal fluorescence micrographs of Caco-2 cells incubated in the absence and presence of polymer treated GFP expressing *V. cholerae* N16961 cultures. *V. cholerae* (green), DNA (blue), and F-actin (red). **C**) Percentage of extracellular LDH in Caco-2 cultures incubated in the absence and presence of polymer treated *V. cholerae* N16961 cultures. Results were normalised to untreated Caco-2 cells (0%) and cells lysed with Triton X-100 (100%). **D**) Levels of cAMP production in Caco-2 cells in the absence and presence of polymer treated *V. cholerae* N16961 cultures. *V. cholerae* cultures were adjusted to an MOI of 10, and incubated in the absence or presence of polymers for 1 h prior to infection of cultured Caco-2 intestinal epithelial cells for 7 h. Results are means ± s.e.m. of three independent experiments. Analysis of variance (ANOVA), followed by Tukey’s post hoc test, was used to test for significance. Statistical significance was defined as p<0.05 (*), p<0.01 (**) or p<0.0001 (****).

While these results suggested that the polymers were able to reduce the pathogenic burden towards Caco-2 cells, with the corresponding decrease in colonisation and toxicity, at this stage, it was unclear if this was a consequence of the sequestration into clusters, or a cumulative effect was being observed, where the repression of virulence, in particular of ctxAB, was contributing to this lower toxicity. Thus, we performed experiments with a quorum sensing mutant (BH1651), that is locked in a low-density phenotype, and thus is unable to switch off the production of CTX toxin at the high-cell densities seen in biofilms.^42,44,45^ Interestingly the effect of clustering on the colonisation of Caco-2 cells by this quorum sensing mutant was very similar to that of the wild type (**Fig. S8**). However, the ability of the polymers to reduce the toxicity of this mutant was significantly compromised (**Fig. S10** and **Fig. S11**), suggesting that the repression of virulence is a significant contributor to the reduced toxicity observed with the wild type. These results were very encouraging, and reflected that sequestration of this pathogen into clusters can result in a reduction of the pathogenic burden, with lower colonisation and toxicity. We thus decided to evaluate the potential of these polymers to prevent colonisation by *V. cholerae* upon ingestion by an animal model. Zebrafish (*Danio rerio*) have been established as an aquatic host which can be colonised and infected by *V. cholerae* in a concentration dependent manner, and infection eventually leads to mortality.^46–48^ Zebrafish are a suitable natural host model for *V. cholerae* colonisation and transmission as their gastrointestinal development and physiology closely mimics that of mammalian organisms.^49^ Additionally, ease of propagation and live imaging made them a good choice of host for our *in vivo* studies, in particular at this early stage. Zebrafish larvae exposed to media containing 10^7^ CFU/mL of GFP-expressing *V. cholerae* for 6 hours were first imaged and then sacrificed, and intestinal *V. cholerae* were extracted from the tissue and enumerated by dilution plating on selective TCBS agar (Fig. 7A). Images of infected fish showed that GFP-expressing *V. cholerae* had specifically colonised the gastrointestinal tract, with the majority of bacteria attached to the mid-intestine (Fig. 7B). Treating the *V. cholerae* contaminated media with polymers significantly reduced the ability of ingested clustered bacteria to colonise zebrafish (~1000-fold) when compared to untreated bacteria (Fig. 7A, top). Ingestion of the residual liquid following removal of the clusters reduced bacterial burdens by only ~100-fold (Fig. 7A, Bassler for sharing strain BH1651. We thank members of the bottom).

**Fig. 7.**
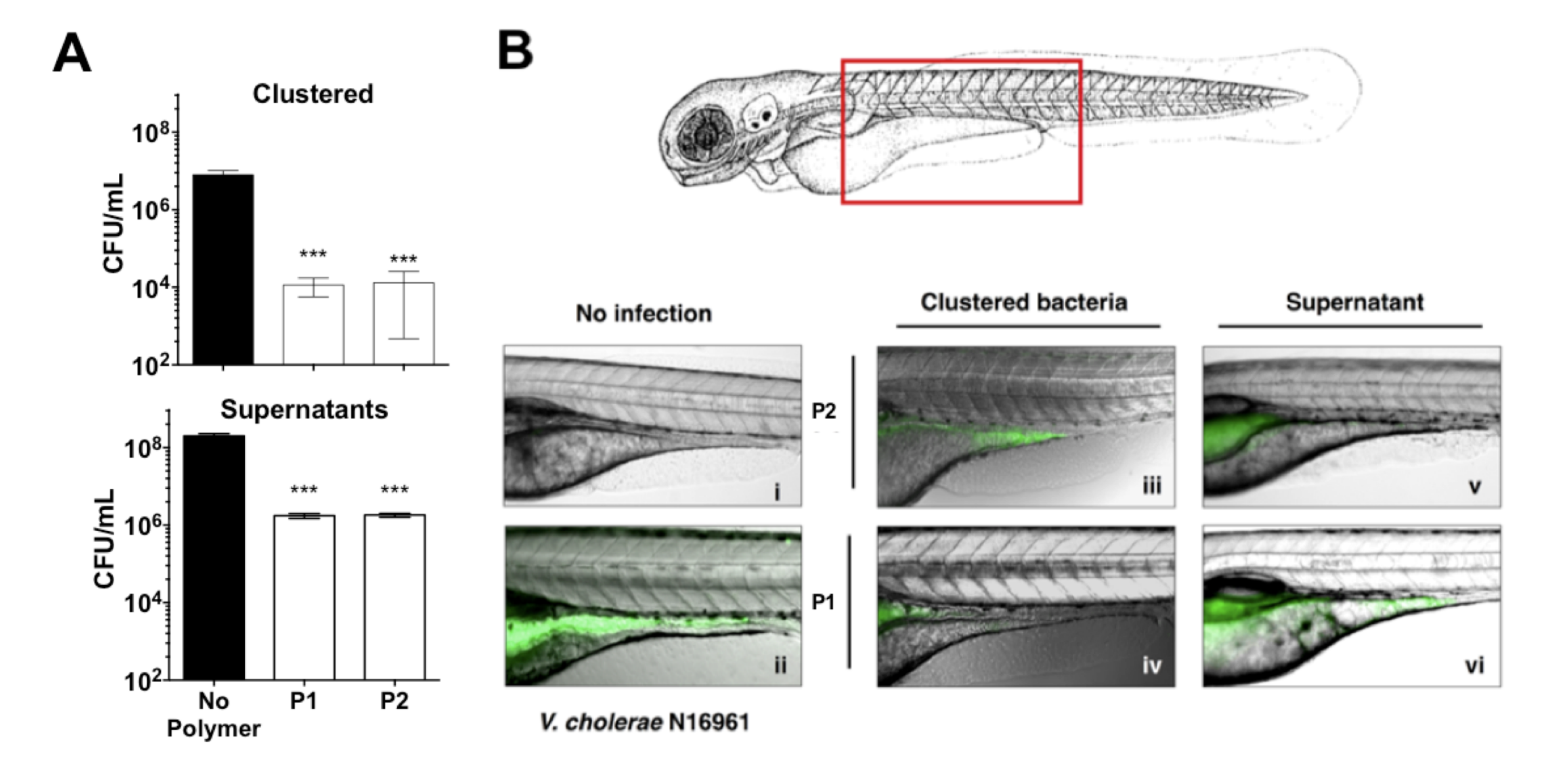
**A)** Number of colony forming units per mL (CFU/mL) of GFP expressing *V. cholerae* N16961 from zebrafish larvae incubated in the absence and presence of polymer treated GFP expressing *V. cholerae* N16961 cultures, following euthanisation and homogenisation of the zebrafish larvae. Zebrafish larvae (n=10 per experimental condition). [**P1**] = 500 μg/mL, [**P2**] = 50 μg/mL. Analysis of variance (ANOVA), followed by Tukey’s post hoc test, was used to test for significance. Statistical significance was defined as p<0.001 (***). B) Overlay of Fluorescence and Optical micrographs of zebrafish larvae left uninfected (panel i), infected with GFP-*V. cholerae* (ii), *V. choler*ae clustered with **P1** (iv) or **P2** (iii), or the remaining decanted supernatant following removal of clustered *V. cholerae* with **P1** (vi) or **P2** (v). [**P1**] = 500 μg/mL, [**P2**] = 50 μg/mL.

## CONCLUSIONS

Here, we have demonstrated that cationic polymers **P1** and **P2**, designed to sequester the human pathogen *V. cholerae* into clusters, can modulate the pathogen’s physiology and induce a transition towards a non-virulent sessile lifestyle. Using a combination of phenotypic and transcriptional assays, we demonstrate that upon clustering, biofilm production is increased, while key virulence factors such as the cholera toxin are repressed. As a result, these polymers can reduce the pathogenic burden in both *in vitro* and *in vivo* models. Sequestration into clusters minimises the ability of the pathogen to colonise both Caco-2 cells and zebrafish, while a reduction in the toxicity of *V. cholerae* to Caco-2 is observed in the presence of the polymers. We anticipate that these findings will be beneficial in the development of novel cost-effective prophylactic or therapeutic polymers, that can engineer microbial responses in unprecedented ways. However, further understanding of the physical and chemical cues that these materials may trigger will need to be developed. Our efforts in this area, as well as in the synthesis of materials with decreased toxicities and higher specificities for the pathogen will be reported in due course.

## Acknowledgements

We thank D. Grainger and his group for their advice on the construction of transcriptional reporter plasmids. We thank B. Bassler for sharing strain BH1651. We thank members of the Krachler and Fernandez-Trillo labs for critical reading and comments on the manuscript. This work was supported by University of Birmingham Fellowships (to A. M. K. and F. F.-T.), Wellcome Trust grant 177ISSFPP (to A. M. K. and F. F.-T), BBSRC grants BB/M021513/1 (to K. V. and A. M. K.) and BB/L007916/1 (to A. M. K.), BBSRC MIBTP studentships (to L. M.) and a CONICYT fellowship (to N. P.-S.).

## AUTHOR CONTRIBUTIONS

All authors contributed to the experimental set-up and discussed the results. A. M. K, F. F.-T. and K. V. secure funding. N. P.-S., L. M., I. I. and D. N. C. synthesised and characterised the polymers. N. P.-S., L. M., K. V., A. M. K. performed the biological assays. N. P.-S., K. V., A. M. K. and F. F.-T. analysed the data, and N. P.-S., A. M. K. and F. F.-T. wrote the manuscript, with all other authors contributing to its final version.

## REFERENCES

1 World Health Organization, Antimicrobial resistance: global report on surveillance, World Health Organization, 2014.

2 European Antimicrobial Resistance Surveillance Network, Antimicrobial resistance surveillance in Europe 2014, 2015.

3 K. Bush, P. Courvalin, G. Dantas, J. Davies, B. Eisenstein, P. Huovinen, G. A. Jacoby, R. Kishony, B. N. Kreiswirth, E. Kutter, S. A. Lerner, S. Levy, K. Lewis, O. Lomovskaya, J. H. Miller, S. Mobashery, L. J. V. Piddock, S. Projan, C. M. Thomas, A. Tomasz, P. M. Tulkens, T. R. Walsh, J. D. Watson, J. Witkowski, W. Witte, G. Wright, P. Yeh and H. I. Zgurskaya, Nat. Rev. Microbiol., 2011, 9, 894–896.

4 C. Nathan, Sci. Transl. Med., 2012, 4, 140sr2.

5 J. O’Neill, The Wellcome TrustHM Government, 2016, 1–84.

6 M.-H. Xiong, Y. Bao, X.-Z. Yang, Y.-H. Zhu and J. Wang, Adv. Drug Delivery Rev., 2014, 78, 63–76.

7 W. Gao, S. Thamphiwatana, P. Angsantikul and L. Zhang, Wiley Interdiscip. Rev.: Nanomed. Nanobiotechnol., 2014, 6, 532–547.

8 A. Som, S. Vemparala, I. Ivanov and G. N. Tew, Biopolymers, 2008, 90, 83–93.

9 R. W. Scott, W. F. Degrado and G. N. Tew, Curr. Opin. Biotechnol., 2008, 19, 620–627.

10 K. Kuroda and G. A. Caputo, Wiley Interdiscip. Rev.: Nanomed. Nanobiotechnol., 2013, 5, 49–66.

11 J. Chen, F. Wang, Q. Liu and J. Du, Chem. Commun., 2014, 50, 14482–14493.

12 A. M. Krachler and K. Orth, Virulence, 2013, 4, 284–294.

13 S. Bhatia, M. Dimde and R. Haag, Med. Chem. Commun., 2014, 5, 862–878.

14 A. Bernardi, J. Jiménez-Barbero, A. Casnati, C. De Castro, T. Darbre, F. Fieschi, J. Finne, H. Funken, K.-E. Jaeger, M. Lahmann, T. K. Lindhorst, M. Marradi, P. Messner, A. Molinaro, P. V. Murphy, C. Nativi, S. Oscarson, S. Penadés, F. Peri, R. J. Pieters, O. Renaudet, J.-L. Reymond, B. Richichi, J. Rojo, F. Sansone, C. Schäffer, W. B. Turnbull, T. Velasco-Torrijos, S. Vidal, S. Vincent, T. Wennekes, H. Zuilhof and A. Imberty, Chem. Soc. Rev., 2013, 42, 4709–4727.

15 E. P. Magennis, A. L. Hook, M. C. Davies, C. Alexander, P. Williams and M. R. Alexander, Acta Biomater, 2016, 34, 84–92.

16 A. H. E. Müller and K. K. Matyjaszewski, Eds., Controlled and Living Polymerizations, Wiley-VCH Verlag GmbH & Co. KGaA, Weinheim, Germany, 2009.

17 World Health Organization, Weekly Epidemiological Record, 2014, 89, 345–356.

18 M. Ali, A. R. Nelson, A. L. Lopez and D. A. Sack, PLOS Negl Trop Dis, 2015, 9, e0003832.

19 M. L. Tamplin, A. L. Gauzens, A. Huq, D. A. Sack and R. R. Colwell, Appl. Environ. Microbiol., 1990, 56, 1977–1980.

20 T. K. Rawlings, G. M. Ruiz and R. R. Colwell, Appl. Environ. Microbiol., 2007, 73, 7926–7933.

21 S. M. Butler and A. Camilli, Nat. Rev. Microbiol., 2005, 3, 611–620.

22 J. K. Teschler, D. Zamorano-Sánchez, A. S. Utada, C. J. A. Warner, G. C. L. Wong, R. G. Linington and F. H. Yildiz, Nat. Rev. Microbiol., 2015, 13, 255–268.

23 A. Saha, A. Rosewell, A. Hayen, C. R. MacIntyre and F. Qadri, Expert Rev Vaccines, 2016, 1–14.

24 E. P. Magennis, F. Fernandez-Trillo, C. Sui, S. G. Spain, D. J. Bradshaw, D. Churchley, G. Mantovani, K. Winzer and C. Alexander, Nat. Mater., 2014, 13, 748–755.

25 A. L. Bole and P. Manesiotis, Adv. Mater. Weinheim, 2016, 28, 5349–5366.

26 X. Xue, G. Pasparakis, N. Halliday, K. Winzer, S. M. Howdle, C. J. Cramphorn, N. R. Cameron, P. M. Gardner, B. G. Davis, F. Fernandez-Trillo and C. Alexander, Angew. Chem., Int. Ed., 2011, 50, 9852–9856.

27 L. T. Lui, X. Xue, C. Sui, A. Brown, D. I. Pritchard, N. Halliday, K. Winzer, S. M. Howdle, F. Fernandez-Trillo, N. Krasnogor and C. Alexander, Nat. Chem., 2013, 5, 1058–1065.

28 I. Louzao, C. Sui, K. Winzer, F. Fernandez-Trillo and C. Alexander, Eur. J. Pharm. Biopharm., 2015, 95, 47–62.

29 E. F. Palermo and K. Kuroda, Biomacromolecules, 2009, 10, 1416–1428.

30 L. C. Paslay, B. A. Abel, T. D. Brown, V. Koul, V. Choudhary, C. L. McCormick and S. E. Morgan, Biomacromolecules, 2012, 13, 2472–2482.

31 J. M. Willey, L. Sherwood and C. J. Woolverton, in Prescotts Principles of Microbiology, Applied Energy, 2008, pp. 126–152.

32 S. M. Faruque, K. Biswas, S. M. N. Udden, Q. S. Ahmad, D. A. Sack, G. B. Nair and J. J. Mekalanos, Proc. Natl. Acad. Sci. U. S. A, 2006, 103, 6350–6355.

33 E. J. Nelson, A. Chowdhury, J. B. Harris, Y. A. Begum, F. Chowdhury, A. I. Khan, R. C. Larocque, A. L. Bishop, E. T. Ryan, A. Camilli, F. Qadri and S. B. Calderwood, Proc. Natl. Acad. Sci. U. S. A., 2007, 104, 19091–19096.

34 C. Reichhardt, J. C. N. Fong, F. Yildiz and L. Cegelski, Biochim. Biophys. Acta, 2015, 1848, 378–383.

35 J. Zhu, M. B. Miller, R. E. Vance, M. Dziejman, B. L. Bassler and J. J. Mekalanos, Proc. Natl. Acad. Sci. U. S. A., 2002, 99, 3129–3134.

36 J. H. Merritt, D. E. Kadouri and G. A. O’Toole, in Current Protocols in Microbiology, John Wiley & Sons, Inc., pp. 1–18.

37 A. K. Dey, A. Bhagat and R. Chowdhury, Journal of Bacteriology, 2013, 195, 2004–2010.

38 K. Skorupski and R. K. Taylor, Mol. Microbiol., 1999, 31, 763–771.

39 J. Sánchez, G. Medina, T. Buhse, J. Holmgren and G. Soberón-Chavez, Journal of Bacteriology, 2004, 186, 1355–1361.

40 B. K. Hammer and B. L. Bassler, Mol. Microbiol., 2003, 50, 101–104.

41 D. A. Higgins, M. E. Pomianek, C. M. Kraml, R. K. Taylor, M. F. Semmelhack and B. L. Bassler, Nature, 2007, 450, 883–886.

42 J. S. Matson, J. H. Withey and V. J. DiRita, Infection and immunity, 2007, 75, 5542–5549.

43 A. J. Silva and J. A. Benitez, PLOS Negl Trop Dis, 2016, 10, e0004330.

44 M. Cámara, A. Hardman, P. Williams and D. Milton, Nat. Genet., 2002, 32, 217–218.

45 W.-L. Ng, L. Perez, J. Cong, M. F. Semmelhack and B. L. Bassler, PLoS Pathog., 2012, 8, e1002767.

46 H. Wang, S. Chen, J. Zhang, F. P. Rothenbacher, T. Jiang, B. Kan, Z. Zhong and J. Zhu, PLoS ONE, 2012, 7, e53383.

47 H. M. Rowe, J. H. Withey and M. N. Neely, Dev. Comp. Immunol., 2014, 46, 96–107.

48 D. L. Runft, K. C. Mitchell, B. H. Abuaita, J. P. Allen, S. Bajer, K. Ginsburg, M. N. Neely and J. H. Withey, Appl. Environ. Microbiol., 2014, 80, 1710–1717.

49 A. N. Y. Ng, T. A. de Jong-Curtain, D. J. Mawdsley, S. J. White, j. Shin, B. Appel, P. D. S. Dong, D. Y. R. Stainier and J. K. Heath, Dev. Biol., 2005, 286, 114–135.

